## Supplemental Figures for "Genome-wide CRISPR interference screen identifies long non-coding RNA loci required for differentiation and pluripotency"

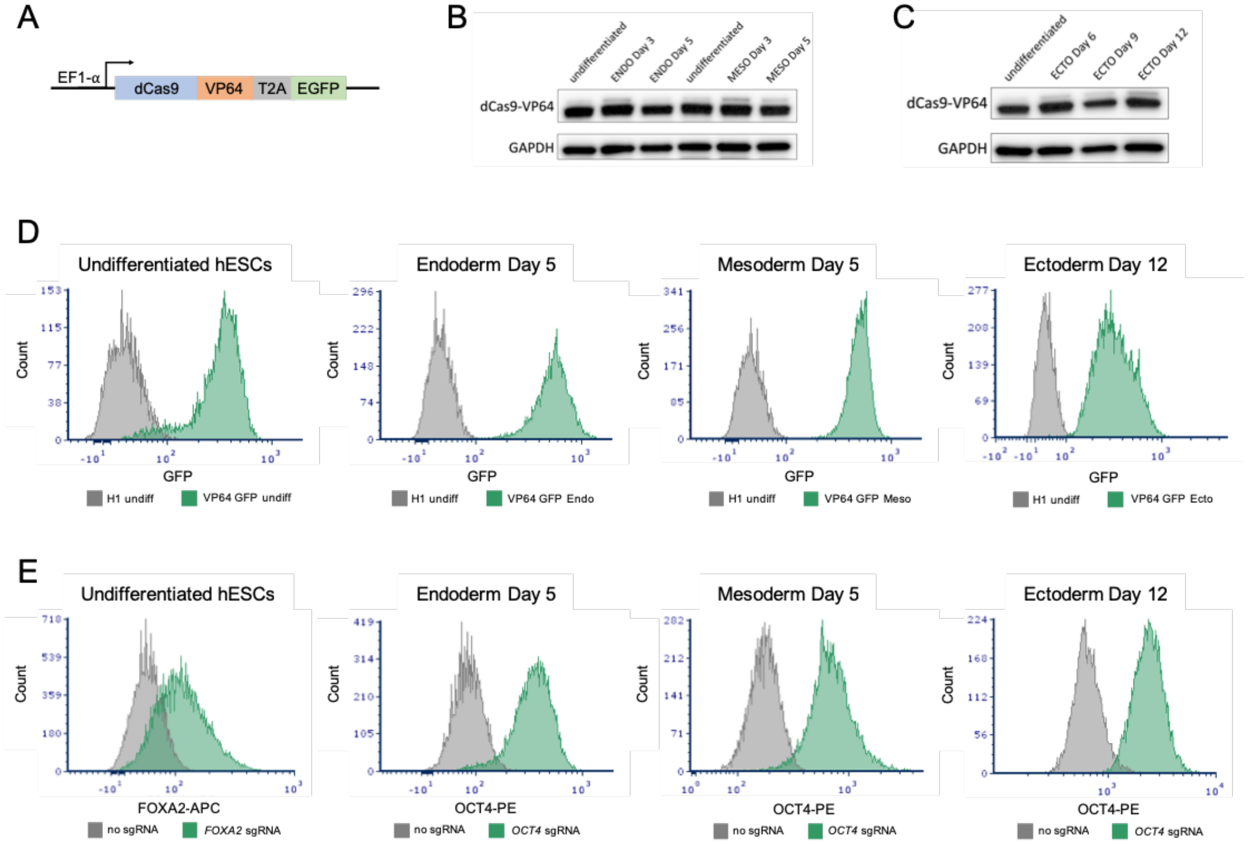


**Figure S1. dCas9-VP64-GFP cell line**
**(A)** Lentiviral GFP construct containing dCas9-VP64 driven by the EF1-α promoter.
**(B)** Western blot of dCas9-VP64 over the course of definitive endoderm and early mesoderm differentiation.
**(C)** Western blot of dCas9-VP64 over the course of neural progenitor cell (ectoderm) differentiation.
**(D)** FACS analysis of GFP expression for undifferentiated and differentiated dCas9-VP64-GFP line compared to H1 cells.
**(E)** FACS staining of targeted mRNA genes in undifferentiated and differentiated dCas9-VP64-GFP cells.

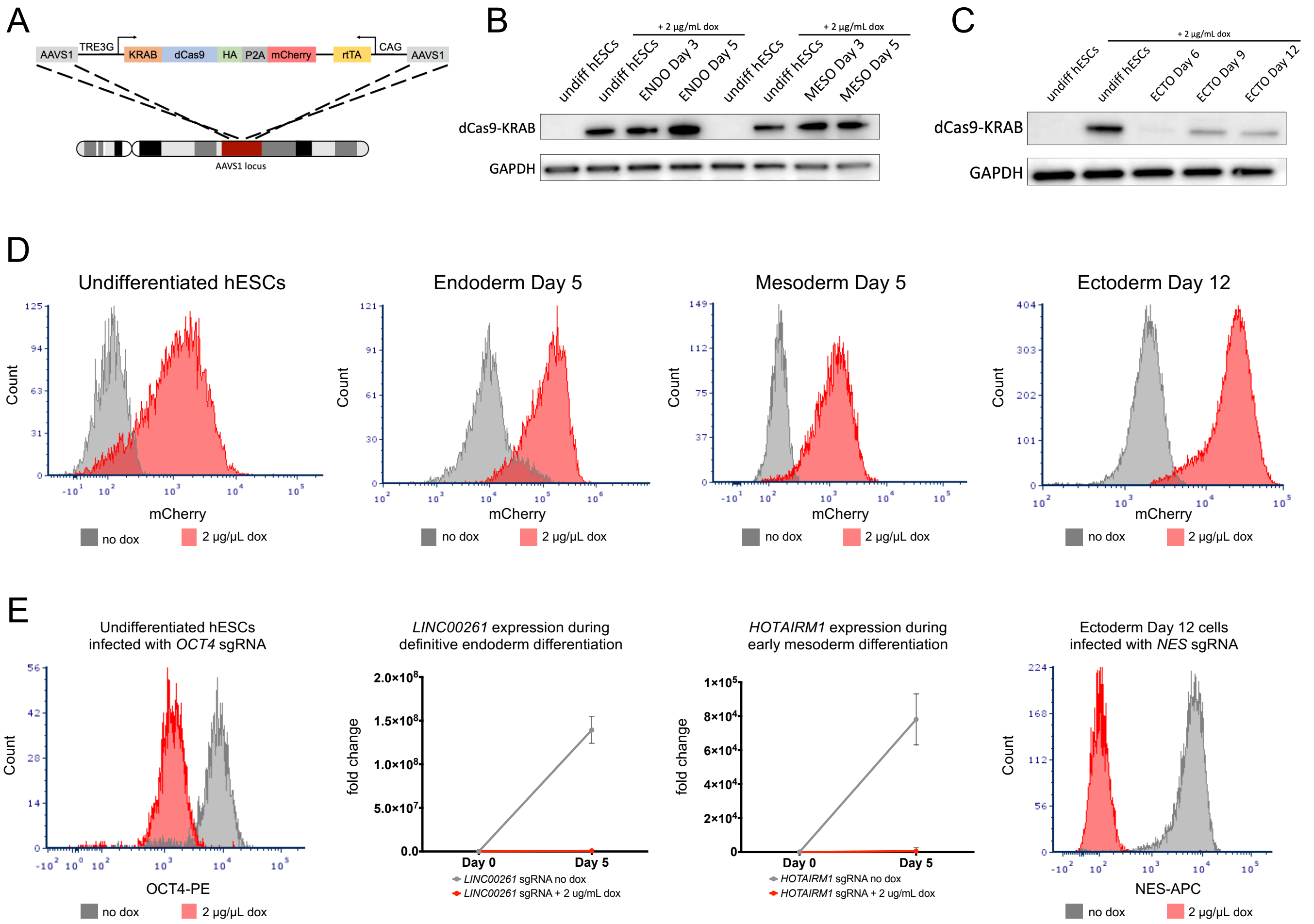


**Figure S2. dCas9-KRAB-mCherry cell line**

**(A)** TALEN construct containing dCas9-KRAB driven by the dox-inducible TRE3G promoter.

**(B)** Western blot of dCas9-KRAB over the course of definitive endoderm and early mesoderm differentiation.

**(C)** Western blot of dCas9-KRAB over the course of neural progenitor cell (ectoderm) differentiation.

**(D)** FACS analysis of mCherry expression for undifferentiated and differentiated dCas9-KRAB-mCherry line compared to “no dox” condition or dCas9-KRAB-GFP line (ectoderm).

**(E)** FACS staining and RT-qPCR expression of targeted mRNA or lncRNA genes, respectively, in undifferentiated and differentiated dCas9-KRAB-mCherry cells.

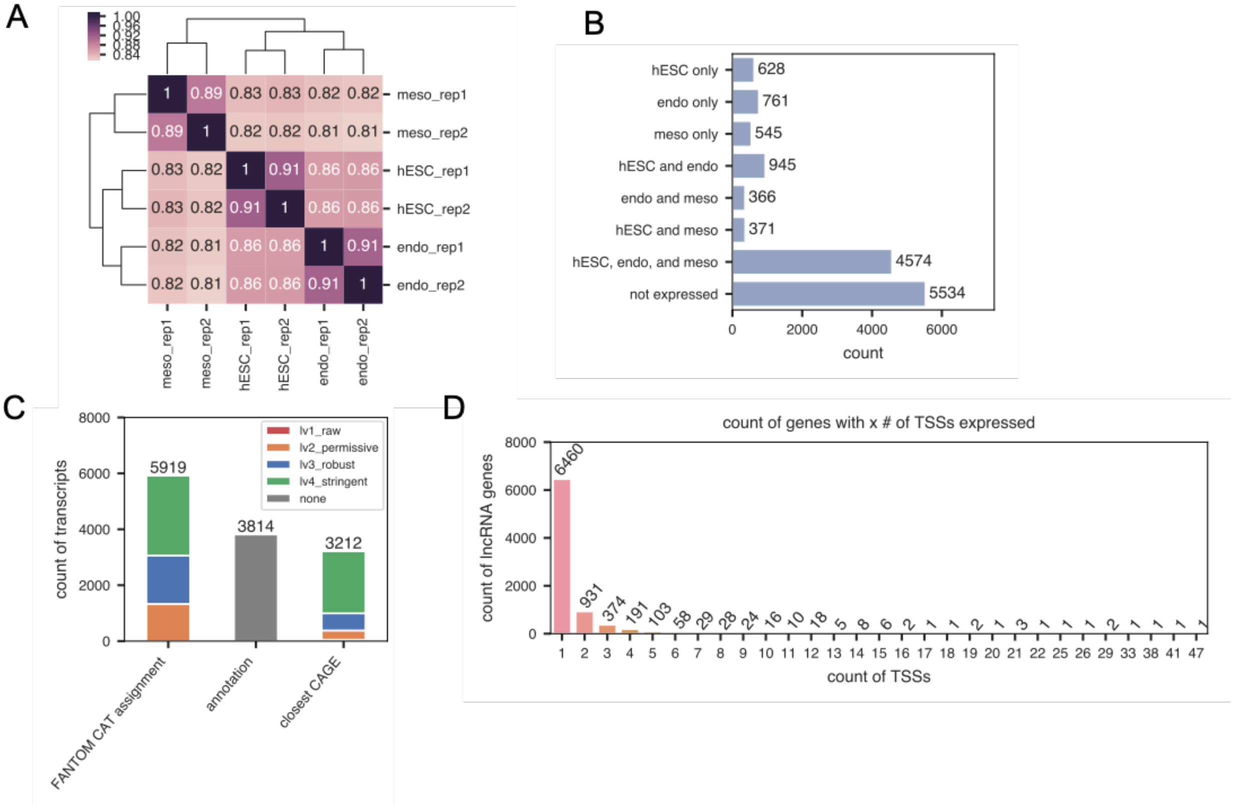


**Figure S3. RNA-sequencing and library design**

**(A)** Spearman correlation of all gene expression values (tpms; both mRNAs and lncRNAs) across biological replicates.

**(B)** Count of lncRNA genes expressed in each subset of lineages, where a lncRNA is counted as expressed if its gene-level tpm is ≥ 0.1 tpm.

**(C)** Count of lncRNA transcripts by TSS assignment strategy: either FANTOM-CAT assignment (Hon et al., 2017), GENCODE TSS annotation, or closest CAGE peak. We only used GENCODE TSS assignments if there was no CAGE peak within 400 bp of the annotated TSS on the same strand. When CAGE peaks were used, the FANTOM5-assigned CAGE peak level is shown in green, blue, orange, and red. The vast majority of assigned CAGE TSSs were defined as either "robust" or "stringent" by FANTOM5.

**(D)** Count of lncRNA genes expressed in any one of the three lineages with a particular number of TSSs assigned; most expressed lncRNAs have only a single assigned TSS.

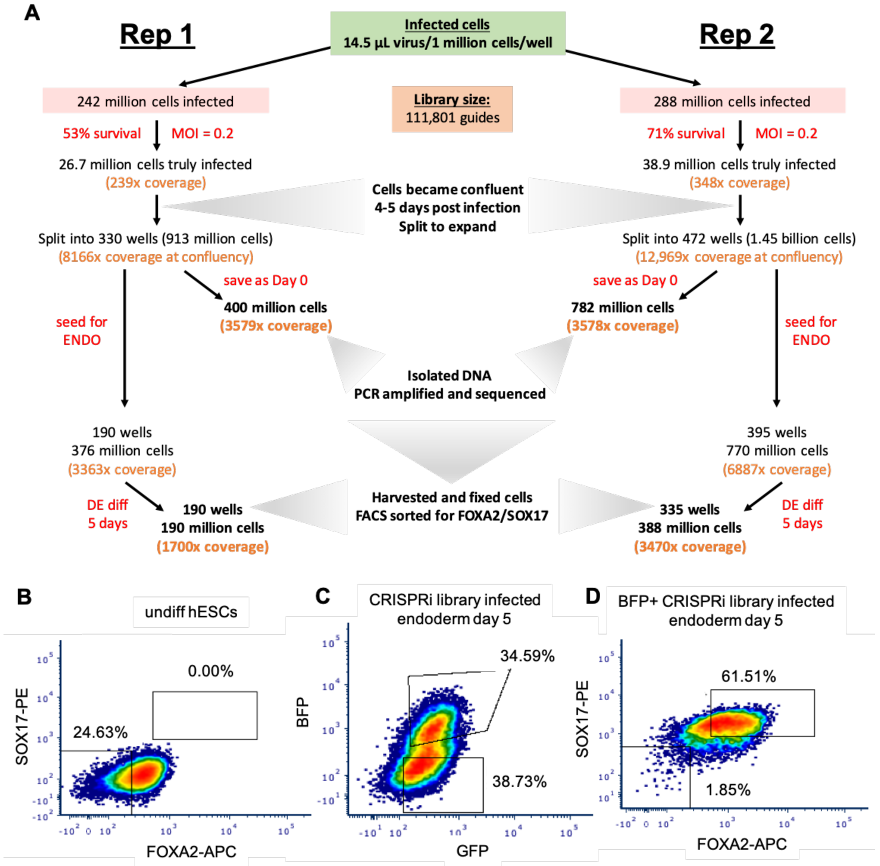


E

| Replicate | Condition | % of cells | # of cells collected |
|---|---|---|---|
| Replicate #1 | BFP+ FOXA2+/SOX17+ | 60% | 37.2 million |
| | BFP+ FOXA2-/SOX17- | 1-2% | 975,000 |
| Replicate #2 | BFP+ FOXA2+/SOX17+ | 60% | 69.4 million |
| | BFP+ FOXA2-/SOX17- | 1-2% | 1.5 million |

**Figure S4. CRISPRi screen schematic and cell numbers**

**(A)** CRISPRi screen schematic. dCas9-KRAB-GFP hESCs (two biological replicates) were infected with pooled sgRNA library, expanded, and seeded out for endoderm differentiation. A population of cells were collected at “Day 0” of endoderm differentiation. Five days post differentiation, cells were harvested, fixed, stained, and sorted for FOXA2/SOX17 double positive and double negative populations. gDNA was subsequently isolated from all populations. The region flanking the sgRNA sequences was PCR amplified and sequenced to determine enrichment of sgRNAs within each population. MOI: multiplicity of infection. DE: definitive endoderm.

**(B)** FACS staining of undifferentiated dCas9-KRAB-GFP hESCs. Cells were fixed and stained with antibodies against FOXA2 and SOX17.

**(C)** BFP/GFP FACS analysis of day 5 endoderm dCas9-KRAB-GFP cells infected with pooled CRISPRi sgRNA library. Cells were stained with FOXA2 and SOX17 antibodies, and subsequently sorted for BFP+ and BFP- conditions (BFP+ marks sgRNA expression).

**(D)** FACS staining of BFP+ sorted cells.

**(E)** Summary of biological replicates of CRISPRi screen.

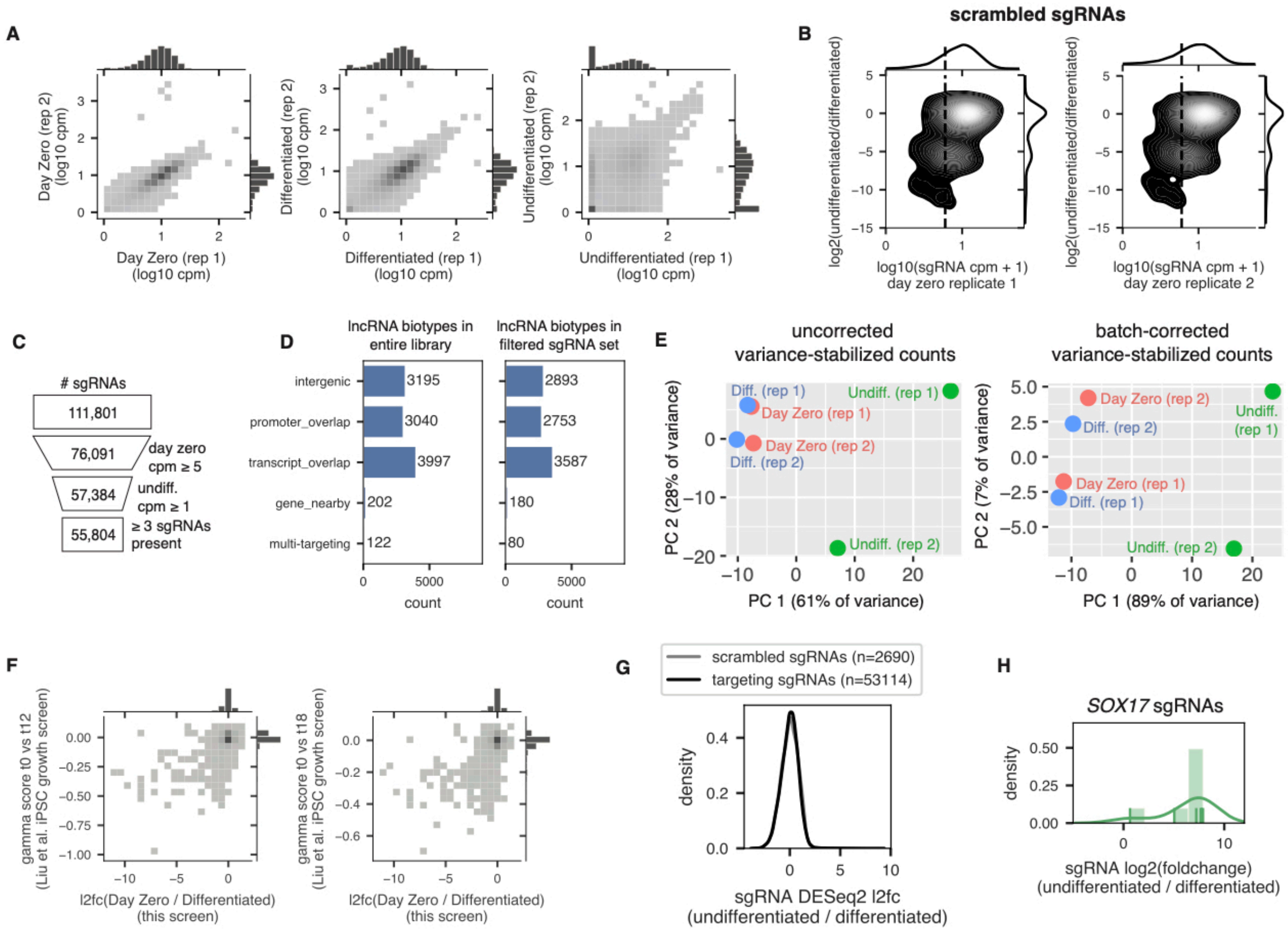

**Figure S5. CRISPRi screen analysis quality control**

**(A)** Bivariate histogram plot showing correspondence of sgRNA counts per million (cpm) for all 111,801 sgRNAs across each set of replicates.

**(B)** Kernel density estimate plot showing the log2 foldchanges of sgRNAs (as calculated by DESeq2) on the y axis compared to the Day Zero cpms on the x axis in log space, for all scrambled sgRNAs in replicate 1 (left) and replicate 2 (right). Vertical line indicates a Day Zero cpm threshold of 5. This threshold eliminates scrambled sgRNAs that show unexpected log2 fold changes due to noise that propagates through the experiment because of low initial counts. Color bar indicates density of sgRNAs.

**(C)** Schematic showing the number of sgRNAs filtered out at each of the 3 filtering steps indicated to result in the final list of 55,804 sgRNAs.

**(D)** Count of lncRNA transcript biotypes represented in the complete library (left) and in the filtered set of sgRNAs (right); biotypes are the same as those outlined in the Figure 2D schematic. Multi-targeting refers to complex TSSs with multiple biotypes within 1000 bp.

**(E)** Left: PCA plot showing the principal components of the DESeq2 variance-stabilized counts (based on the 100 sgRNAs with the highest variance across samples); right: analogous PCA plot after removing batch effects across replicates (using limma's removeBatchEffect function). Note that the batch-corrected data is shown in the PCA for illustration purposes only; in the actual calculation of log2 fold changes, raw, unmodified counts were used and batch (replicate) was simply added to the DESeq2 model as a covariate.

**(F)** Bivariate histogram plot showing correspondence of sgRNA drop-out between our screen (x axis) and the CRiNCL iPSC growth screen (y axis) for 1,117 sgRNAs that overlapped between the two. For our screen, drop out was calculated using DESeq2 (log2 fold change between Day Zero and differentiated replicates). Shown are two of the drop out scores calculated by the CRiNCL authors (left: time 0 vs. time 12, right: time 0 vs. time 18).

**(G)** Density plot showing the DESeq2 log2 foldchanges of scrambled sgRNAs (gray) and targeting sgRNAs (black); the two densities overlap almost perfectly.

**(H)** Density plot showing the DESeq2 log2 foldchanges of sgRNAs targeting *SOX17,* where individual sgRNAs are highlighted by vertical lines on the x axis.

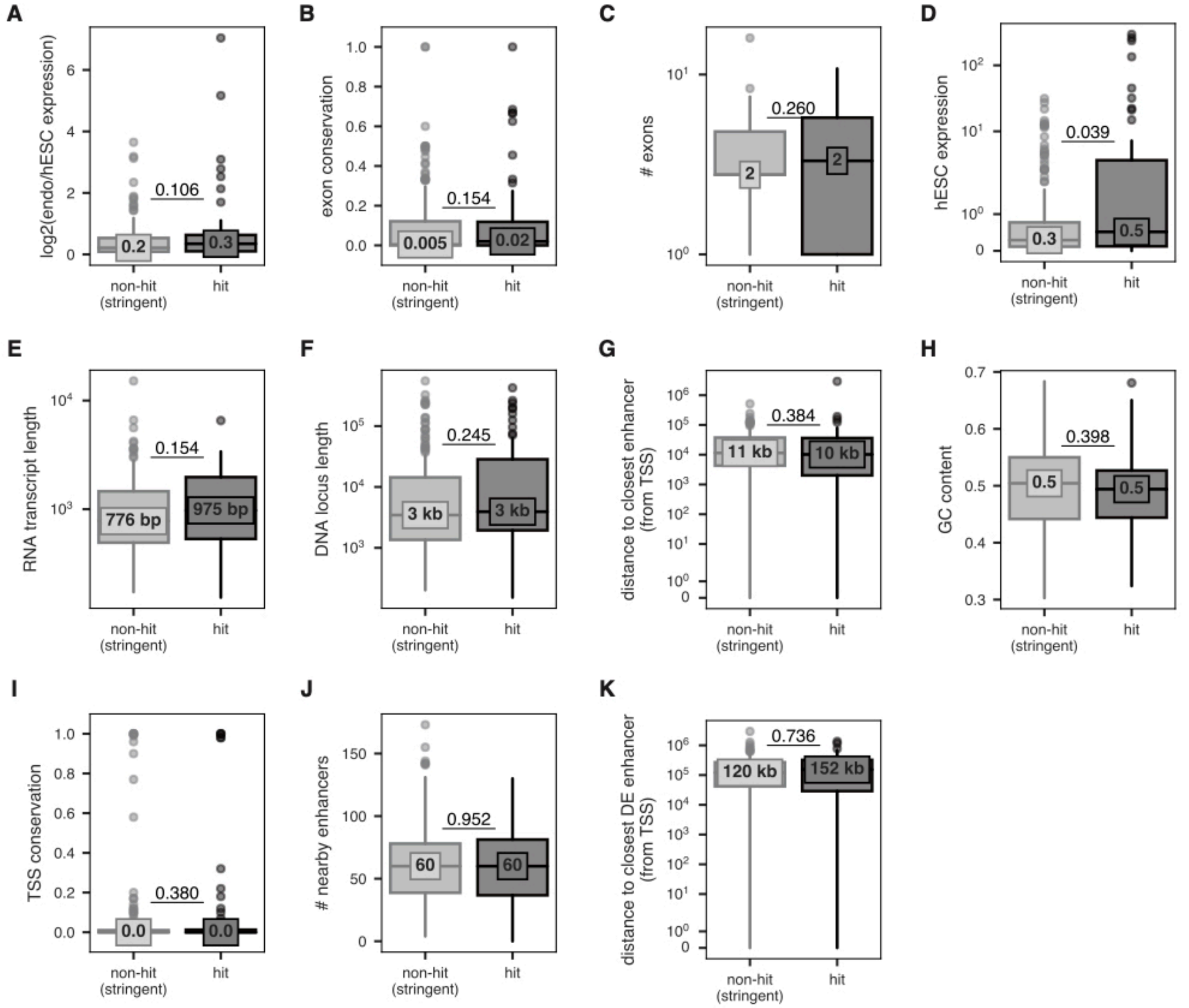


**Figure S6. Common features of lncRNA hits**

Additional features compared between hit lncRNA loci and stringent non hit lncRNA loci that are not significant at an alpha of < 0.05 by a two-sided Mann Whitney test. Exact, uncorrected p-values are listed, and median of each distribution is highlighted.

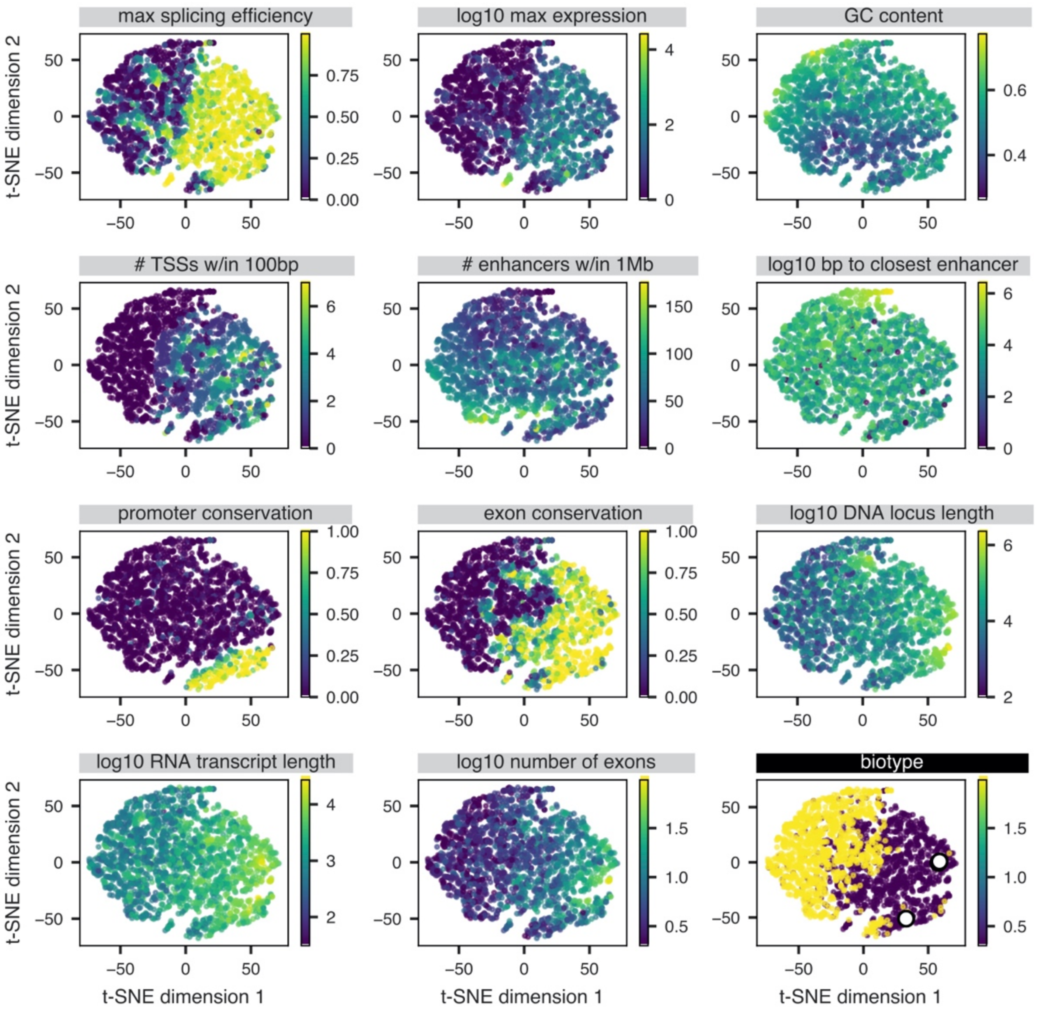


**Figure S7. t-SNE visualization of 11 genomic features**
Visualization of the 11 genomic features outlined in **Figure 6A** using the same t-SNE as in **Figures 6B-D**. In each subplot, genes are colored by a given feature. In the final subplot showing the genes colored by biotype, the white dots indicate the 3 gold standard lncRNAs used in **Figure 6C** (*XIST, NEAT1,* and *MALAT1*). In all plots, a subsample of the data is shown to facilitate plotting (random sample of 1000 lncRNAs and random sample of 1000 mRNAs; the same random sample that is shown in **Figure 6**).

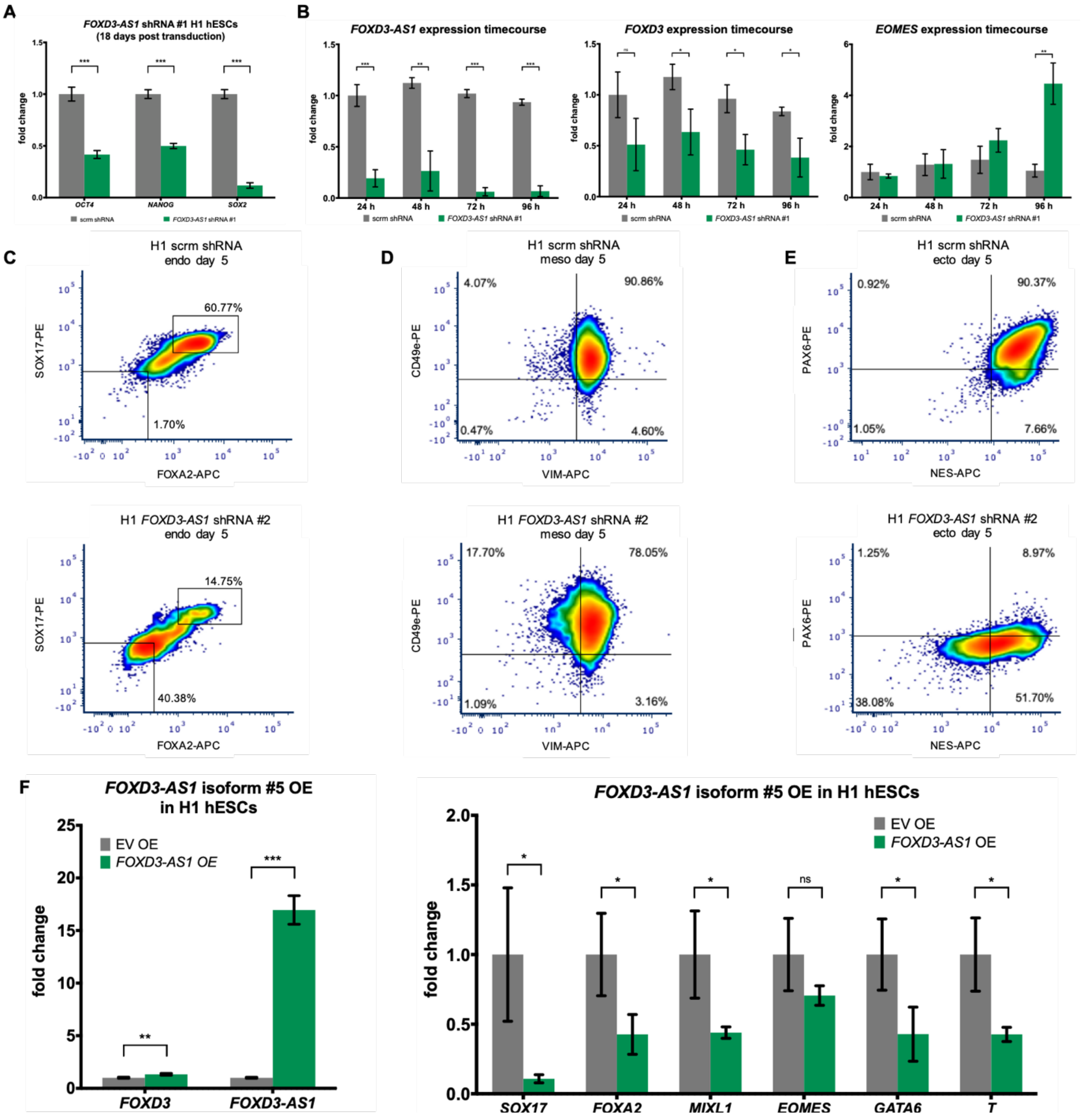


**Figure S8. *FOXD3-AS1* knockdown and overexpression**

**(A)** RT-qPCR expression of pluripotency marker genes 18 days post infection of H1 hESCs with *FOXD3-AS1* shRNA. *** = $p < 0.001$ by an unpaired t-test.

**(B)** RT-qPCR expression time course during *FOXD3-AS1* shRNA infection of H1 hESCs. * = $p < 0.05$, ** = $p < 0.01$, *** = $p < 0.001$ by an unpaired t-test.

**(C-E)** FACS staining of day 5 definitive endoderm cells (C), day 5 early mesoderm cells (D), or day 12 neural progenitor cells (E) infected with scrambled shRNA or FOXD3-AS1 shRNA #2. Cells were fixed and stained with antibodies against FOXA2 and SOX17 (C), VIM and CD49e (D), or NES and PAX6 (E).

**(F)** RT-qPCR expression of *FOXD3-AS1* and endoderm/mesoderm genes 9 days post infection of H1 hESCs with *FOXD3-AS1* isoform #5 expression vector. *** = $p < 0.001$ by an unpaired t-test.

| lncRNA hit gene | nearby gene TSS | nearby gene type | nearby gene known endo regulator? |
|---|---|---|---|
| AC006077.3 | PCBD2 | protein_coding | Yes |
| AC011523.2 | KLK15 | protein_coding | No |
| AC022201.5 | FAM136A | protein_coding | No |
| AC068831.3 | UNC45A | protein_coding | No |
| AC093627.9 | AC093627.10 | lncRNA | No |
| ACVR2B-AS1 | ACVR2B | protein_coding | Yes |
| CTC-525D6.1 | VSTM2B | protein_coding | No |
| CTD-2308L22.1 | EPHA4 | protein_coding | Yes |
| CTD-2545G14.4 | DLG4 | protein_coding | No |
| CTD-2631K10.1 | TNPO1 | protein_coding | No |
| DIGIT | GSC | protein_coding | Yes |
| FOXD3-AS1 | FOXD3 | protein_coding | Yes |
| HOXC-AS1 | HOXC9 | protein_coding | Yes |
| LAMTOR5-AS1 | LAMTOR5 | protein_coding | No |
| LINC01424 | UBE2G2 | protein_coding | No |
| MKLN1-AS | MKLN1 | protein_coding | No |
| PCBP1-AS1 | PCBP1 | protein_coding | No |
| PITPNA-AS1 | INPP5K | protein_coding | No |
| RAMP2-AS1 | RAMP2 | protein_coding | No |
| RP1-292L20.3 | PCNX1 | protein_coding | No |
| RP11-120D5.1 | HCCS | protein_coding | Yes |
| RP11-121L10 | CHORDC1 | protein_coding | No |
| RP11-259N19.1 | HK2 | protein_coding | No |
| RP11-326C3.12 | IFITM3 | protein_coding | Yes |
| RP11-326I11.3 | IRF2 | protein_coding | No |
| RP11-359E3.4 | BMPR1A | protein_coding | Yes |
| RP11-402J6.1 | NEUROG2 | protein_coding | Yes |
| RP11-421L21.3 | DPH5 | protein_coding | No |
| RP11-615I2.2 | CHD3 | protein_coding | No |
| RP11-867G23.8 | B4GAT1 | protein_coding | No |
| RP3-508I15.9 | TOMM22 | protein_coding | No |
| RP4-621N11.2 | RBL1 | protein_coding | No |
| RP4-680D5.8 | DNAJC16 | protein_coding | No |
| RP5-1065J22.8 | TMEM167B | protein_coding | No |

**Figure S9. lncRNA hits with other nearby TSSs**
The 35 lncRNA hits with other nearby TSSs (within 1000 bp of the lncRNA's TSS) are shown, along with the gene name of the nearby TSS, its biotype (lncRNA or protein-coding), and whether the nearby gene is known to be a regulator of hESC differentiation, as determined via a literature search (those that are near known regulators are colored in green).

### SUPPLEMENTAL TABLE DESCRIPTIONS

**Supplemental Table S1. RNA-seq lncRNA expression in hESCs, endoderm, and mesoderm, related to Figure 2**

**Supplemental Table S2. CRISPRi screen sgRNA counts in each population, related to Figure 3**

**Supplemental Table S3. CRISPhieRmix results per targeted transcript in screen, related to Figure 3**

**Supplemental Table S4. Validation results for the 22 sgRNAs tested individually, related to Figure 4**

**Supplemental Table S5. Genomic features of all lncRNAs targeted in the screen, related to Figure 5**

**Supplemental Table S6. List of cancers/traits from endoderm-derived tissues, used for GWAS SNPs analysis, related to Figure 5**

**Supplemental Table S7. Clustering results of all annotated lncRNAs and mRNAs, related to Figure 6**
